# Insulin B peptide-MHC class II-specific chimeric antigen receptor-Tregs prevent autoimmune diabetes

**DOI:** 10.1101/2023.02.23.529737

**Authors:** Justin A. Spanier, Vivian Fung, Christine M. Wardell, Mohannad H. Alkhatib, Yixin Chen, Linnea A. Swanson, Alexander J. Dwyer, Matthew E. Weno, Nubia Silva, Jason S. Mitchell, Paul C. Orban, Majid Mojibian, C. Bruce Verchere, Brian T. Fife, Megan K. Levings

**Author notes:** **Co-Corresponding Authors:** Brian T. Fife, 2101 6th St SE, Wallin Medical Biosciences Building, 3-146, Minneapolis, MN 55455., Megan K. Levings, A4-186, 950 West 28^th^ Ave, Vancouver, BC, Canada. Equal contribution (Justin A. Spanier, Vivian Fung).

## Abstract

Adoptive immunotherapy with Tregs is a promising approach for prevention or treatment of type 1 diabetes. Islet antigen-specific Tregs have more potent therapeutic effects than polyclonal cells, but their low frequency is a barrier for clinical application. To generate Tregs that recognize islet antigens, we engineered a chimeric antigen receptor (CAR) derived from a monoclonal antibody with specificity for the insulin B-chain 10-23 peptide presented in the context of the IA^g7^ MHC class II allele present in NOD mice. Peptide specificity of the resulting InsB-g7 CAR was confirmed by tetramer staining and T cell proliferation in response to recombinant or islet-derived peptide. The InsB-g7 CAR re-directed NOD Treg specificity such that insulin B 10-23-peptide stimulation enhanced suppressive function, measured via reduction of proliferation and IL-2 production by BDC2.5 T cells and CD80 and CD86 expression on dendritic cells. Co-transfer of InsB-g7 CAR Tregs prevented adoptive transfer diabetes by BDC2.5 T cells in immunodeficient NOD mice. In wild type NOD mice, InsB-g7 CAR Tregs stably expressed Foxp3 and prevented spontaneous diabetes. These results show that engineering Treg specificity for islet antigens using a T cell receptor-like CAR is a promising new therapeutic approach for the prevention of autoimmune diabetes.

**Brief Summary:** Chimeric antigen receptor Tregs specific for an insulin B-chain peptide presented by MHC class II prevent autoimmune diabetes.

## Introduction

Autoimmune type 1 diabetes (T1D) is caused by T cell mediated destruction of the insulin producing beta cells of the islets of Langerhans in the pancreas. Regulatory T cells (Tregs), defined by expression of the Foxp3 transcription factor, are suppressive CD4^+^ T cells that normally function to limit autoreactive effector T cell responses and prevent autoimmunity (1-3). Tregs from individuals with T1D have abnormal cytokine and gene expression profiles, and reduced suppressive function, leading to the concept that strategies to restore Treg function could be a promising way to treat or prevent autoimmunity (4-8).

Significant research using animal models of T1D has shown that therapeutic restoration of Tregs can prevent disease progression. Clinical studies have shown the safety of this approach in humans, but so far evidence for efficacy is limited (9-11). A consideration is that, to date, all clinical trials of Treg therapy in T1D have used polyclonal cells, meaning that only a small fraction of the infused cells were specific for disease-relevant antigens (12). In animal models of autoimmune diabetes, there is clear evidence that islet-antigen-specific Tregs are significantly more effective than polyclonal Tregs at preventing or delaying disease. In animal models of autoimmune diabetes, BDC2.5 TCR-transgenic Tregs, which are specific for a fusion peptide between insulin C and chromogranin A, or NOD T cells engineered to express Foxp3 and the BDC2.5 T cell receptor (TCR), but not polyclonal Tregs, suppressed diabetes induced by T cells from diabetic NOD mice or BDC2.5 T effector cells (13, 14). Similarly, in wild type (WT) NOD mice, a single infusion of BDC2.5 TCR-transgenic Tregs prevented and reversed spontaneous T1D, whereas polyclonal cells had no effect (15, 16).

In addition to engineering Tregs with TCRs, an alternate method to re-direct specificity is through use of chimeric antigen receptors (CARs), engineered receptors in which antigen-specificity is typically mediated by an extracellular single chain antibody (scFv) and intracellular signaling driven by one or more co-stimulatory domains and CD3ζ. In comparison to TCRs, advantages of using CARs to control T cell specificity include the high affinity and specificity of scFvs, modular intracellular domain format, lack of mispairing with endogenous TCR chains, and MHC independence. We and others have shown that CAR-Tregs specific for allo or autoantigens have potent therapeutic effects in models of allograft rejection or autoimmunity (17-24). A limitation of CARs is that they are most effective when cross-linked by membrane or oligomeric proteins, possibly explaining why Tregs expressing a CAR specific for soluble insulin did not prevent or delay T1D in the NOD spontaneous diabetes model (25).

Seeking to design CARs for use in Tregs to treat T1D, we considered that the immunosuppressive function of Tregs needs to be present in both affected tissues and local lymph nodes (26). We thus hypothesized that the ideal CAR-Treg target in T1D would be a protein with expression restricted to islets and also present within the pancreas-draining lymph nodes. Therefore, we developed a new mAb specific for the amino acid (AA) residues 10-23 of the insulin B chain (InsB) presented in the context of NOD mouse MHCII, IA^g7^. We then converted this mAb to a CAR and tested its ability to re-direct Treg specificity and create a therapeutic cell product that could suppress diabetogenic T cells and prevent autoimmune diabetes.

## Results

### Generation and validation of an InsB_10-23_ :IA^g7^-specific mAb

Our goal was to design a CAR that enhanced retention of, and suppression by, Tregs at the site of islet-antigen presentation. Therefore, we first generated mAbs with high specificity for an islet-antigen peptide presented by NOD MHCII, IA^g7^. Specifically, we targeted insulin B chain AA residues 10-23 because transgenic and retrogenic CD4^+^ T cells targeting this epitope are capable of causing autoimmune diabetes and CD4^+^ T cell responses to this epitope are required for spontaneous disease in NOD mice (27-30). To generate an InsB_10-23_:IA^g7^ mAb, B cell hybridomas were created from NOD mice immunized with recombinant IA^g7^ molecules containing InsB_10-23_ altered peptides InsBP8E and InsBP8G (31, 32) (Figure 1A). These modified insulin peptides were used because of their high affinity and well-defined binding to IA^g7^. In contrast, the native WT InsB_10-23_ binds weakly to IA^g7^, is unstable, and is presented in multiple registers (33-35). One of the resulting hybridomas, named 1B2, produced an IgG1 antibody that specifically bound to IA^g7^ tetramers containing InsB_10-23_ modified peptides, but not to IA^g7^ tetramers containing irrelevant peptides human CLIP_87-101_ or hen egg lysozyme (HEL)_11-25_ (Figure 1B). To further confirm specificity, we used fluorochrome conjugated 1B2 to stain peptide-pulsed bone marrow-derived dendritic cells (BMDC). BMDCs pulsed with InsBP8E, and to a lesser extent P8G, showed a significant increase in 1B2 geometric mean fluorescent intensity compared to BMDCs pulsed with vehicle control (Figure 1, C and D). Using biolayer interferometry we confirmed a preference for InsBP8E over InsBP8G, where the relative affinity (K_D_) of 1B2 for InsBP8E:IA^g7^ was nearly 3-fold that of InsBP8G:IA^g7^ (8.5×10^−9^M vs 2.7×10^−8^M, respectively).

**Figure 1.**
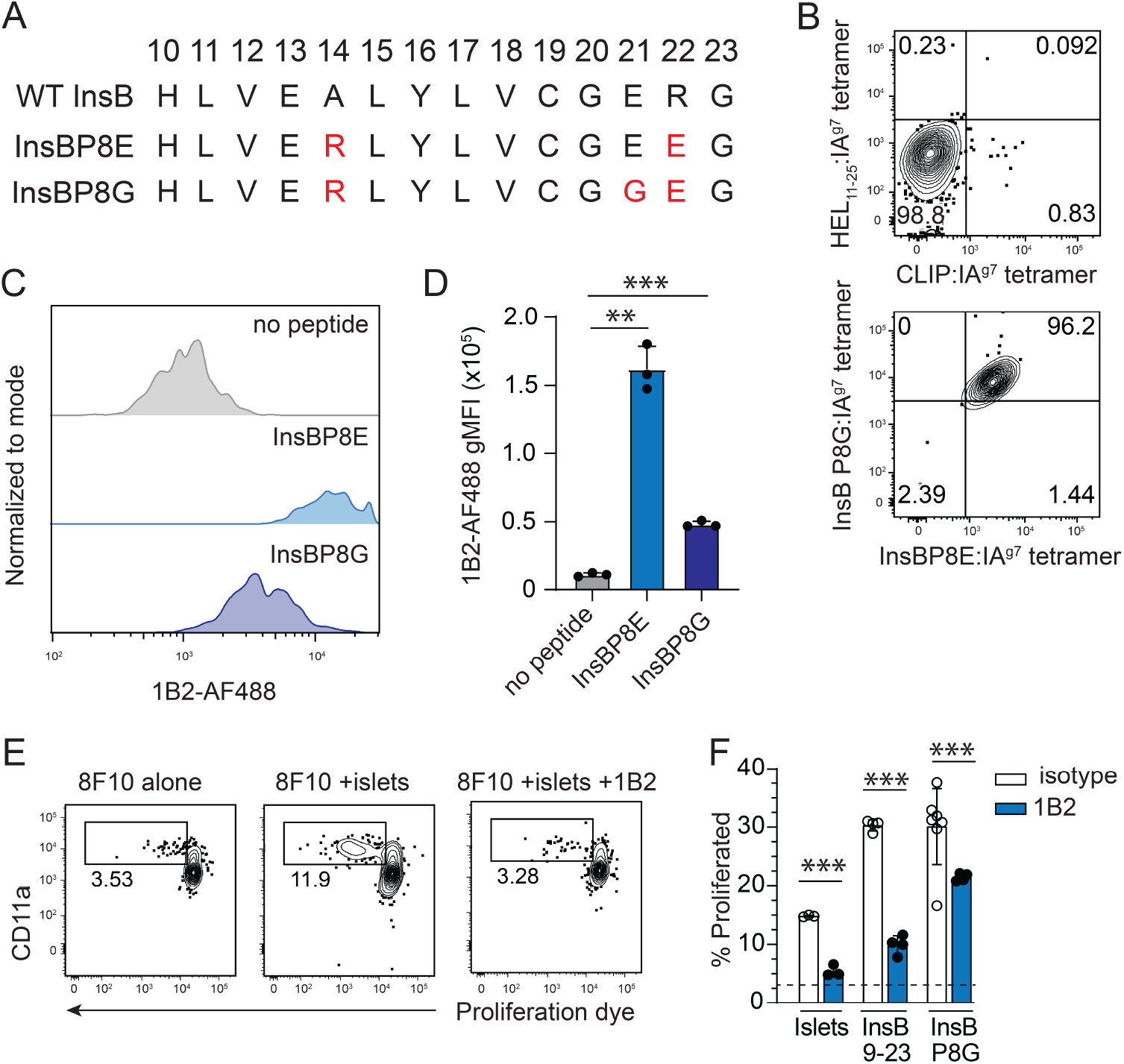
Generation and validation of an InsB10-23:IAg7-specific mAb. (**A**) Amino acid sequence of wild type (WT) InsB_10-23_ compared to its mimotopes InsBP8E and InsBP8G. Position of each amino acid within the insulin B chain peptide is shown on top. Residues highlighted in red font differ from the WT peptide sequence. (**B**) Flow cytometry plots showing staining of the 1B2 hybridoma with the indicated tetramer. (**C**) Flow cytometry histograms showing AF488-labeled 1B2 antibody staining of BMDCs pulsed overnight with the indicated peptide. (**D**) Quantification of the data shown in (**C**). Data are representative of 4 independent experiments. (**E**) Representative flow cytometry plots of 8F10 TCR transgenic CD4^+^ T cells labeled with cell trace violet and stimulated *in vitro* for 4 days with APCs and islets in the presence or absence of 50μg 1B2 blocking antibody. (**F**) Quantification of (**E**), comparing the frequency of unstimulated 8F10 T cells to those stimulated with InsB_10-23_, InsBP8G, or NOD islets in the presence of 50μg blocking 1B2 or IgG1 isotype control antibody. The dashed line represents the average proliferation of 8F10 T cells in the absence of antigen and mAb.Data are pooled from two independent experiments, n=3-9/group. One-way ANOVA with multiple comparisons, \*\*\**P*<0.001.

To further characterize antibody specificity we next determined if the 1B2 antibody could block CD4^+^ T cell responses to endogenous, islet-derived antigen. InsB_10-23_-specific TCR-transgenic 8F10 T cells were cocultured with NOD splenocytes pulsed with islets or peptides in the presence of 1B2 or IgG1 isotype control antibody (28). 8F10 CD4^+^ T cells proliferated in response to both InsB_10-23_ and InsBP8G peptides and to a lesser extent, NOD islets (Figure 1, E and F). Addition of 1B2 antibody significantly reduced 8F10 CD4^+^ T cell proliferation in response to all three stimuli (InsB_10-23_, InsBP8G, and islets).

In another test of 1B2 specificity, we used the AS150 T cell hybridoma, which is specific for InsB_10-23_ peptide and secretes IL-2 in response to InsBP8E pulsed antigen presenting cells (APCs) (33, 36). At low peptide concentrations, this response was inhibited with addition of 1B2 antibody (Supplemental Figure 1). In contrast, 1B2 had no effect on BDC2.5 CD4^+^ T cell proliferation in response to the cognate antigen 2.5 hybrid-peptide (2.5HP), a fusion peptide composed of Insulin C and chromogranin A (Supplemental Figure 2). These data demonstrate that the 1B2 antibody is specific for the InsB_10-23:_IA^g7^ peptide-MHC complex. Importantly, its affinity is sufficient to inhibit cognate T cell activation and proliferation in the presence of naturally processed and presented InsB_10-23_ antigen.

**Figure 2.**
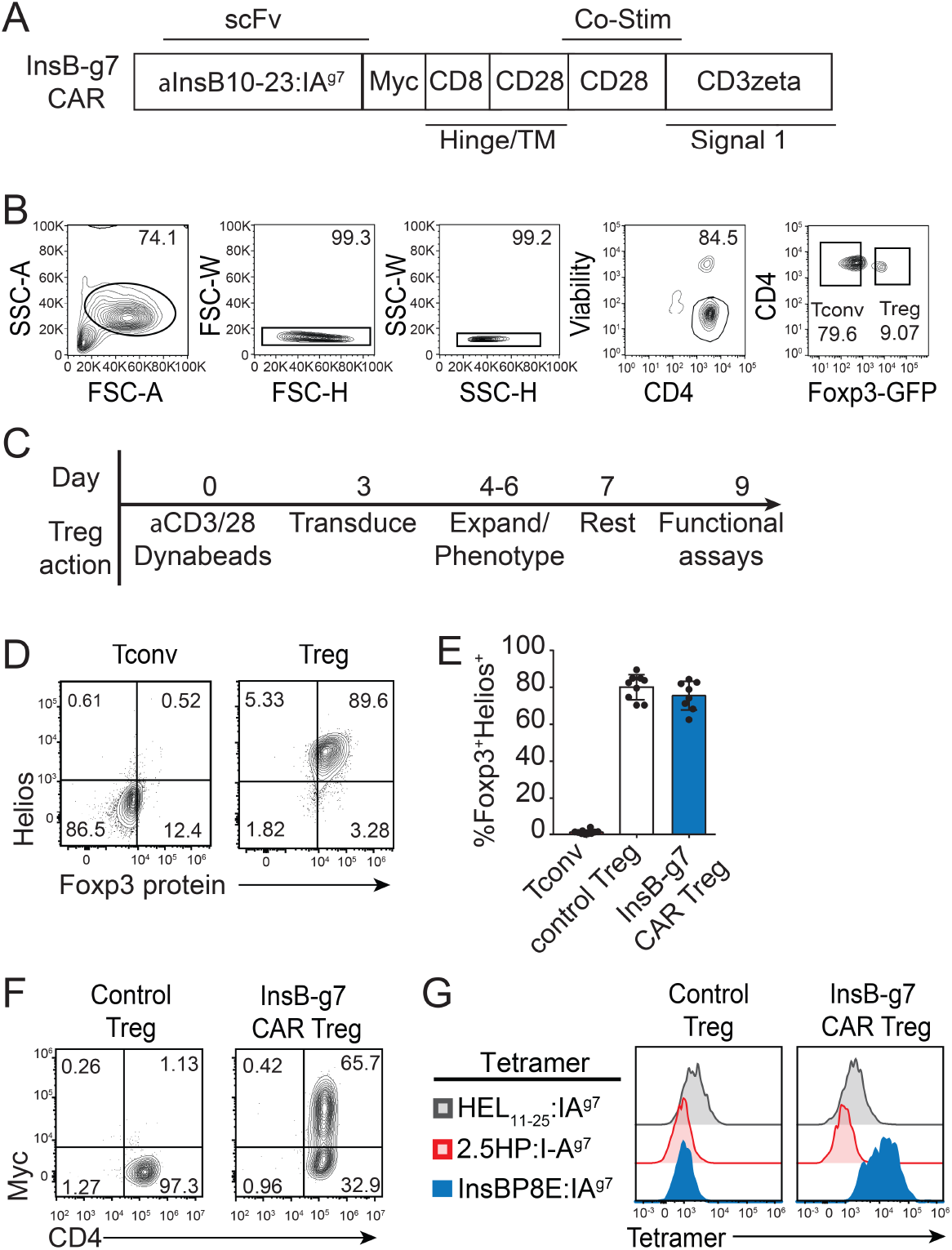
*In vitro* expanded InsB-g7 CAR Treg cells retain Foxp3 expression and bind cognate antigen. (**A**) Schematic depicting the design of the InsB-g7 CAR derived from the variable region of the 1B2 antibody. (**B**) Representative flow cytometry plots showing gating strategy used to sort GFP^+^ Tregs from NOD.Foxp3^EGFP^ mice. (**C**) Timeline of protocol used to engineer and expand CAR Tregs from NOD mice. (**D**) Representative flow cytometry plots showing purity of sorted and expanded GFP^+^ Treg compared to GFP^−^ Tconv cells. Cells are gated on size, viability, CD4^+^ T cells. (**E**) Quantification of the purity of control Tregs and InsB-g7 CAR Treg cells compared to expanded GFP^−^ Tconv cells. Data are pooled from 8-9 experiments with each data point representing one individual experiment. (**F**) Representative flow cytometry plots showing CAR expression, as assessed by Myc tag staining, in InsB-g7 CAR Tregs compared to control Tregs. (**G**) Tetramer staining of Myc^+^ InsB-g7 CAR Tregs compared to Myc^−^ control Tregs. Data are representative of 9 experiments.

### 1B2 CAR Tregs are stable and retain specificity for InsB_10-23_

We next converted the 1B2 mAb into a CAR by linking heavy and light chain variable domain sequences from the 1B2 antibody and cloning upstream of an extracellular Myc-epitope tag followed by CD28 and CD3ζ intracellular signaling domains (17) (Figure 2A). To characterize the specificity of the resulting InsB-g7 CAR, Tregs were sorted from NOD.Foxp3^EGFP^ reporter mice and transduced with retrovirus encoding the InsB-g7 CAR or control (Figure 2, B and C). Compared to GFP^neg^ Tconv cells, sorted CAR Treg cells were >90% Foxp3^+^, and the majority co-expressed Helios, another transcription factor characteristic of the Treg lineage (Figure 2, D and E). Furthermore, InsB-g7 CAR Tregs bound the InsBP8E:IA^g7^ tetramer but not the 2.5HP:IA^g7^ tetramer (Figure 2, F and G). Collectively, these data demonstrate that when reformatted as a CAR, the 1B2 antibody scFv had preserved insulin peptide-MHC complex specificity, and that NOD Treg specificity could be re-directed towards an islet antigen upon expression of this CAR.

### InsB-g7 CAR T cells are activated by synthetic and naturally-presented insulin peptides

To further test the specificity and function of the InsB-g7 CAR, we assessed the proliferative response of InsB-g7 CAR T cells to InsB_10-23_ peptide. 1B2 CAR Tconvs or Tregs were labeled with a proliferation dye and cocultured *in vitro* with irradiated NOD splenocytes, in the presence of an irrelevant peptide control (HEL_11-25_) or InsB_10-23_ P8E. In response to InsBP8E peptide both InsB-g7 CAR Tconvs and Tregs underwent significant proliferation (Figure 3, A-D). Consistent with the hypo-proliferative nature of Tregs (37), InsB-g7 CAR Tconvs proliferated more extensively in response to InsBP8E than InsB-g7 CAR Tregs, but the cumulative division index (CDI) was greater in InsB-g7 CAR Tregs due to lower background proliferation in the absence of exogenous antigen.

**Figure 3.**
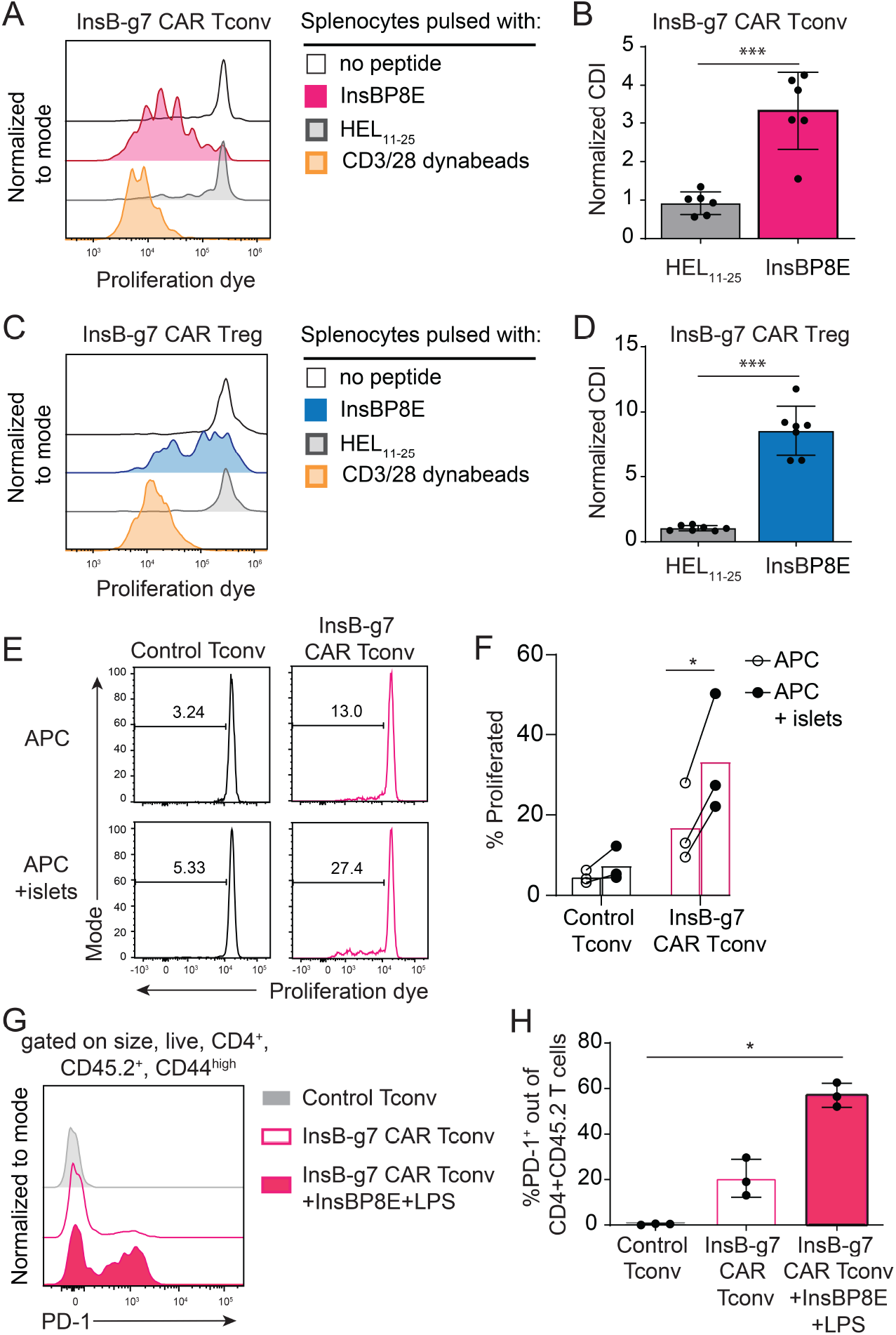
InsB-g7 CAR T cells are activated by synthetic and naturally presented insulin peptides. (**A**) Representative flow cytometry histograms showing *in vitro* proliferation of InsB-g7 CAR Tconv following 3 day coculture with splenocytes pulsed with no peptide, InsB_10-23_, HEL_11-25_, or anti-CD3/CD28 antibody coated dynabeads. (**B**) Cell division index (CDI)-based quantification of (**A**). Data are from 6 independent experiments. One-way ANOVA with multiple comparisons, \*\*\**P*<0.001. (**C**) Representative flow cytometry plots showing *in vitro* proliferation of InsB-g7 CAR Tregs following 3 day coculture with splenocytes pulsed with no peptide, InsB_10-23_, HEL_11-25_, or CD3/CD28 coated dynabeads. (**D**) Quantification of (**C**). Data are from 7 independent experiments. One-way ANOVA with multiple comparisons, \*\*\**P*<0.001. (**E**) Representative flow cytometry histograms showing control Tconv and InsB-g7 CAR Tconv cell proliferation after 3 day coculture with APCs alone or together with NOD islets. (**F**) Quantification of (**E**). Data are from 3 experiments. Two-way ANOVA with multiple comparisons, \**P*<0.05. (**G**) Representative flow cytometry plots showing PD-1 expression on control Tconv and InsB-g7 CAR Tconv cells isolated from the spleen 7 days post-transfer into 8 week old female NOD mice. One of the treatment groups received InsBP8E peptide with LPS intravenously on day 0. (**H**) Quantification of (**G**). Data are representative of 2 independent experiments, n=3 per group. One way ANOVA with multiple comparisons, \**P*<0.05.

We next tested if the InsB-g7 CAR could stimulate T cell activation in response to islet-derived antigen. These experiments were done with Tconv due to their higher *in vitro* proliferative capacity compared to Tregs. Control Tconvs or InsB-g7 CAR Tconvs were labeled with a proliferation dye and cocultured *in vitro* with T cell-depleted NOD splenocytes in the presence or absence of NOD islets. A significant fraction of InsB-g7 CAR Tconvs (∼30%) proliferated in response to islets, whereas control Tconvs did not (Figure 3, E and F).

To test if InsB-g7 CAR Tconvs could respond to endogenous islet-derived antigen *in vivo*, control or InsB-g7 CAR Tconvs were injected into 8 week old female NOD mice and 7 days later the T cells were isolated from the secondary lymphoid organs and analyzed by flow cytometry for PD-1 expression, as an indirect measure of stimulation (38). Whereas control Tconvs had no detectable PD-1 expression, ∼20% of InsB-g7 CAR Tconvs expressed PD-1. As a positive control, mice were immunized with InsBP8E and lipopolysaccharide (LPS), resulting in nearly 60% of 1B2 CAR Tconvs staining positive for PD-1 (Figure 3, G and H). These data show that the InsB-g7 CAR can detect islet-derived antigen *in vivo* and, similar to what is observed with TCR ligation, become activated and upregulate PD-1.

### InsB-g7 CAR Tregs mediate bystander suppression in vitro

After confirming that InsB-g7 CAR Tregs were stimulated by islet-derived antigen, we next assessed their antigen-stimulated suppressive capacity. We first tested if CAR stimulation induced expression of proteins related to Treg activation and suppression. Control Tregs and InsB-g7 CAR Tregs were cocultured *in vitro* overnight in the presence of NOD splenocytes with either InsBP8E, HEL_11-25_, or no peptide. We found that InsBP8E stimulation led to a significant increase in CD69, LAP, and CTLA4 expression on InsB-g7 CAR Tregs compared to HEL_11-25_-stimulated InsB-g7 CAR Tregs; no effect was observed in control Tregs, regardless of peptide antigen (Figure 4A). Another *in vitro* correlate of *in vivo* suppressive function is antigen-stimulated transendocytosis of costimulatory molecules CD80 and CD86 from APCs (39). When InsB-g7 CAR Tregs were cocultured with splenic CD11c^+^ dendritic cells (DCs) pulsed with InsBP8E, both CD80 and CD86 expression on DCs was significantly reduced compared to DCs pulsed with HEL_11-25_ (Figure 4, B and C). In contrast, there was no change in CD80/86 expression on DCs pulsed with InsBP8E in the presence of control Tregs (Figure 4, B and C). These data show that stimulation of InsB-g7 CAR Tregs with cognate antigen results in increased expression of molecules associated with Treg suppression and facilitates enhanced transendocytosis of CD80 and CD86 from APCs.

**Figure 4.**
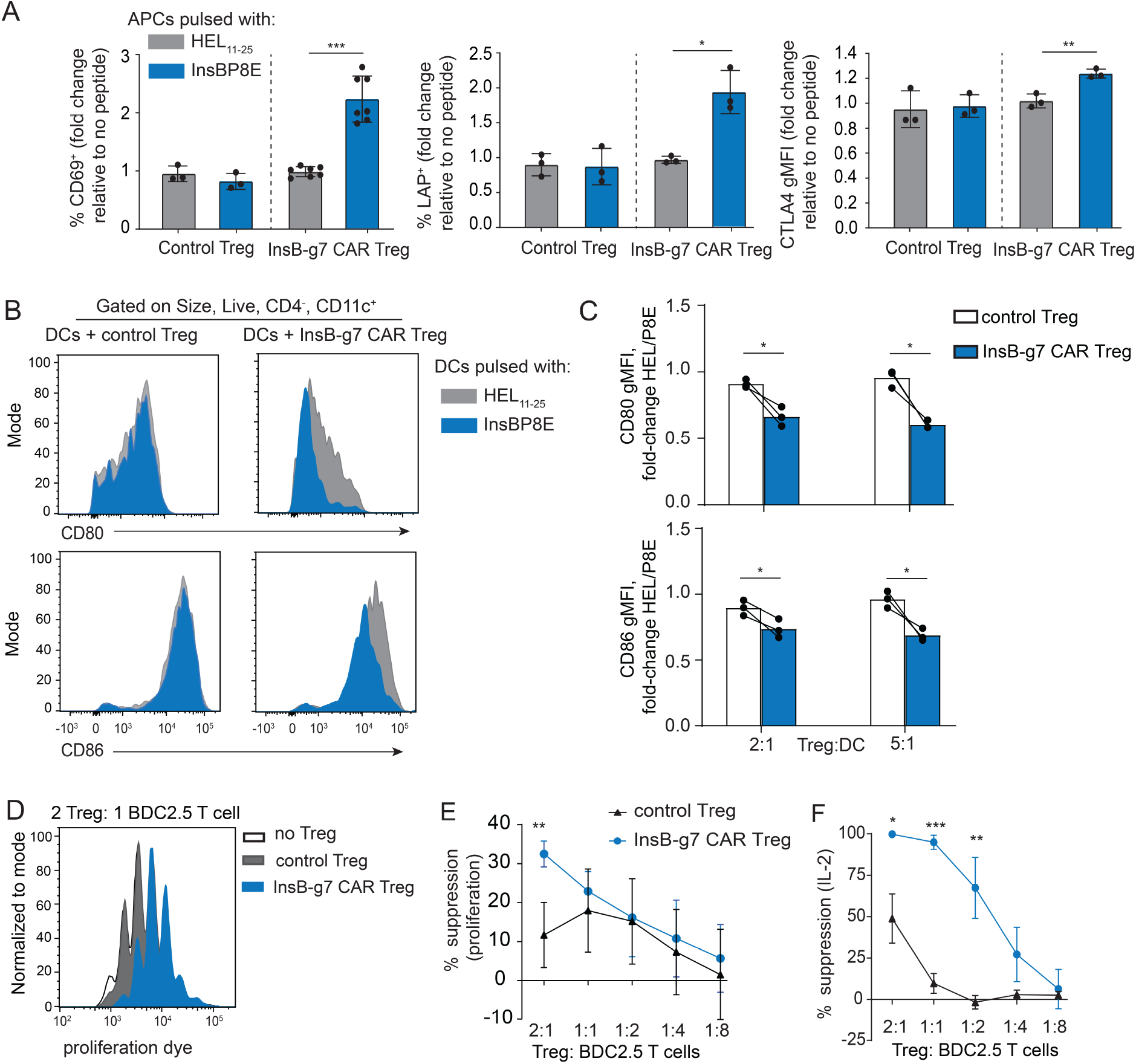
InsB-g7 CAR Tregs mediate bystander suppression in vitro. (**A**) Quantification of flow cytometry data showing the fold-change in the frequency of Tregs expressing CD69 (left) and LAP (middle), and gMFI of CTLA-4 (right) expression on InsB-g7 CAR Tregs or control Tregs after overnight coculture with splenocytes pulsed with InsBP8E or HEL_11-25_, relative to cells cocultured in the absence of peptide. Data are from at least three independent experiments, n=3-7/group. One-way ANOVA with multiple comparisons, \**P*<0.05, \*\**P*<0.01, \*\*\**P*<0.001. (**B**) Representative histograms showing CD80 (top) and CD86 (bottom) expression on HEL_11-25_ or InsB_10-23_ pulsed splenic CD11c+ DCs following 2 day coculture with control Treg or InsB-g7 Treg at 2:1 Treg:DC ratio. (**C**) Quantification of (**B**) represented as fold-change in CD80 (top) or CD86 (bottom) gMFI of DCs pulsed with InsBP8E relative to HEL_11-25_. Data are from 3 independent experiments. Paired t-test, \**P*<0.05. (**D**) Representative flow cytometry data from *in vitro* Treg suppression assays showing proliferation of BDC2.5 T cells in the presence of antigen loaded DCs and control or InsB-g7 CAR Tregs. (**E**) Quantification of (**D**). (**F**) Quantification of IL-2 cytokine in the culture supernatants from (**D**). Data in (**E**) and (**F**) are normalized to the no Treg group and are from 2 independent experiments, n=3/group Two-way ANOVA with multiple comparisons of control Treg group to InsB-g7 CAR Treg group at each Treg:T cell ratio, \**P*<0.05,\*\**P*<0.01, \*\*\**P*<0.001.

We next determined if CAR stimulation enhanced Treg suppression *in vitro* using BDC2.5 T cells as responders. InsB-g7 CAR or control Tregs were cocultured overnight with DCs in the presence of the peptides InsBP8E and P63, a mimotope for the BDC2.5 TCR. The next day, CD4^+^ BDC2.5 T cells were added to Treg/DC cultures and BDC2.5 T cell proliferation was assessed by flow cytometry after 3 additional days. Compared to the no Treg control, both control Tregs and InsB-g7 CAR Tregs suppressed BDC2.5 T cell proliferation and IL-2 production (Figure 4, D-F). However, at the 2:1 Treg:T cell ratio, the InsB-g7 CAR Tregs were significantly more suppressive than control Tregs. Moreover, InsB-g7 CAR Tregs suppressed IL-2 production significantly more than control Tregs at multiple ratios (Figure 4, D-F). Collectively, these data show InsB-g7 CAR stimulation leads to increased expression of markers of activation and suppression, resulting in the suppression of DCs and effector function of T cells with disparate antigen specificity.

### InsB-g7 CAR Tregs suppress BDC 2.5 T cell-induced autoimmune diabetes

To test the *in vivo* function of InsB-g7 CAR Tregs we first used an adoptive transfer model of autoimmune diabetes. In this model the Teff:Treg ratio can be controlled and antigen encounter synchronized. We transferred 50×10^3^ naïve BDC2.5 CD4^+^ T cells alone or together with either control Tregs or InsB-g7 CAR Tregs at 3:1 or 9:1 Treg:BDC2.5 T cell ratio into NOD.RAG^KO^ recipient mice (Figure 5A). Prior to adoptive transfer, Tregs expressed high levels of Foxp3 compared to sorted, GFP^neg^ Tconv cells, and >90% of InsB-g7 CAR Tregs bound to InsBP8G tetramer, demonstrating high Treg purity and CAR expression (Supplemental Figure 3). Mice that were given BDC2.5 T cells alone developed diabetes within 14 days, whereas mice co-injected with InsB-g7 CAR Tregs at either 9:1 or 3:1 Treg:BDC2.5 T cell ratios were completely protected from autoimmune disease (Figure 5B). In contrast, 3/9 mice and 2/4 mice co-injected with control Tregs at 9:1 and 3:1 Treg:BDC2.5 T cell ratios, respectively, developed diabetes with kinetics similar to mice treated with BDC2.5 T cells alone.

**Figure 5.**
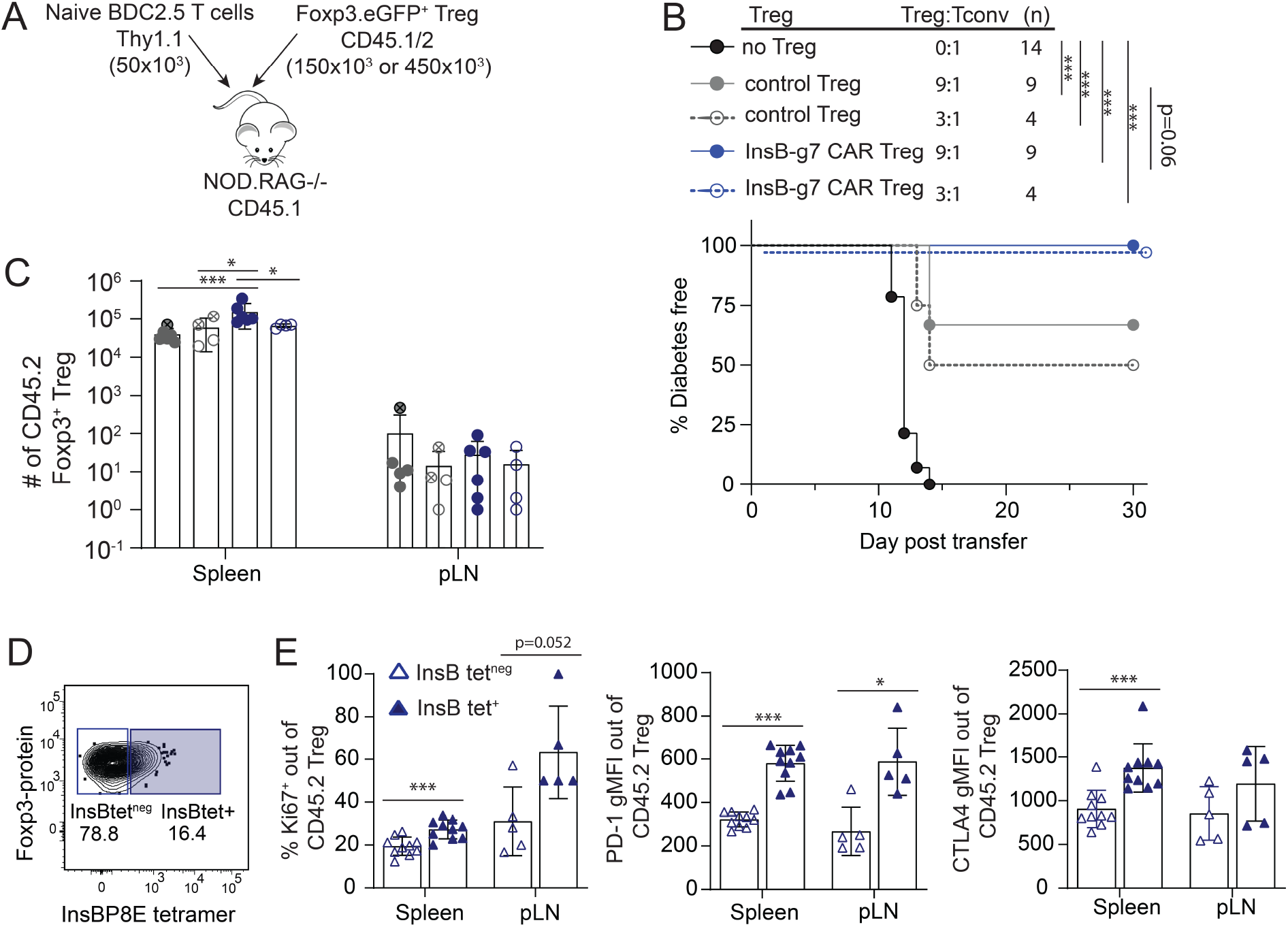
InsB-g7 CAR Tregs suppress BDC 2.5 T cell-induced autoimmune diabetes. (**A**) Experimental design indicating that 450×10^3^ or 150×10^3^ control or InsB-g7 CAR Tregs were co-transferred with 50×10^3^ naïve BDC2.5 CD4^+^ T cells into 6-10wk old NOD.RAG^KO^ recipient mice. (**B**) Cell ratio and diabetes free survival of mice described in (**A**). Data are from 2-3 independent experiments, n=4-14 mice/group. Log-rank test with Bonferroni correction, \*\*\**P*<0.001. (**C**) Quantification of CD4^+^CD45.2^+^Foxp3^+^ Tregs isolated from spleen and pLN from mice treated with 9:1 or 3:1 Treg:Tconv, as indicated in (**B**). Circles containing crosshairs depict diabetic mice, with all mice analyzed either 2 days after becoming diabetic or at day 30 post-transfer. (**D**) Representative flow cytometry plot illustrating gating strategy for identification of CD4^+^CD45.2^+^ InsBP8E tetramer^+^ and tetramer^neg^ Foxp3^+^ Tregs. (**E**) The frequency of Ki67^+^ and gMFI intensity of surface PD-1 and intracellular CTLA4 of tetramer^neg^ and tetramer^+^ InsBg7 CAR Tregs. Data are pooled from mice treated at 9:1 and 3:1 ratios in 2-3 independent experiments, n=5-10mice/group. Only mice with >10 CD4^+^CD45.2^+^ T cells (limit of detection) were included in the analysis. Student’s t-test, \**P*<0.05, \*\*\**P*<0.001.

To gain more insight into how InsB-g7 CAR Tregs suppressed autoimmune diabetes we used flow cytometry to enumerate and phenotype the CAR Tregs and BDC2.5 T cells either within two days of disease onset or, for those mice that remained nondiabetic, 30 days post cell transfer. Using CD45.2 as a congenic marker to identify the CAR Tregs, we found that the spleens of mice receiving the InsB-g7 CAR Tregs at a 9:1 ratio (Treg:BDC2.5) contained significantly more CD45.2^+^ Tregs than those of mice receiving InsB-g7CAR Tregs at a 3:1 ratio or mice that received control Treg (Figure 5C). We next determined if protection from autoimmune disease in InsB-g7 CAR Treg treated mice was associated with enhanced InsB-g7 CAR Tregs proliferation and suppressive phenotype. While nearly all CD45.2^+^ cells retained Foxp3 expression, only a subset maintained surface expression of the InsB-g7 CAR, as assessed by InsBP8E tetramer staining (Figure 5D). Comparing the phenotype of InsBP8E tetramer^+^ and tetramer^neg^ cells, the tetramer^+^ cells showed higher expression of Ki67 in both the spleen and pLN (Figure 5E). Furthermore, the tetramer^+^ InsB-g7 CAR Tregs also showed significantly higher levels of PD-1 and CTLA4 (Figure 5E), showing that InsB-g7 CAR Tregs were stimulated *in vivo*. Thus, InsB-g7 CAR Treg treated mice had the highest numbers of CAR Tregs in the spleen and the lowest incidence of autoimmune diabetes.

### InsB-g7 CAR Tregs reduce the number of BDC 2.5 T effector cells in peripheral lymphoid organs and pancreas

We next examined the effects of InsB-g7 CAR Tregs on BDC2.5 T cell expansion and/or effector cytokine production. All Treg-treated mice had significantly reduced numbers of BDC2.5 T cells in the spleen, whereas mice treated with InsB-g7 CAR Tregs at 9:1 Treg:Tconv ratio also had a significantly reduced number of BDC2.5 T cells in the pLN (Figure 6A). In addition to reduced BDC2.5 T cell numbers, InsB-g7 CAR Treg treatment led to a significant reduction in the number of IFNγ- and TNFα-producing polyfunctional BDC2.5 T effector cells in the spleen (Figure 6B). While all mice receiving Tregs had a significant reduction in the BDC Tconv:Treg ratio, mice receiving the highest dose of InsB-g7 CAR Tregs had the lowest ratio of BDC Tconv:Treg (Figure 6B). Using immunofluorescent staining of pancreas tissue either at the time of diabetes diagnosis or 30 days post transfer, we found Foxp3^+^ InsB-g7 CAR Tregs in close islet proximity, in peri-insulitis and within islets (Figure 6D). Histologically there was a significant difference in the amount of BDC2.5 T cell insulitis in the pancreas of mice treated with InsB-g7 CAR Tregs, despite no significant difference in disease incidence between Treg treated groups (p=0.06), (Figure 6, D and E). At both the 9:1 and 3:1 Treg:BDC2.5 T cell ratios, mice treated with InsB-g7 CAR Tregs showed a significant, nearly 10-fold reduction in the total area around the islets that was occupied by BDC2.5 Tconv cells (9:1 p<0.01, 3:1 p<0.05). In summary, InsB-g7 CAR Tregs likely prevent autoimmunity via suppression of BDC2.5 T effector function.

**Figure 6.**
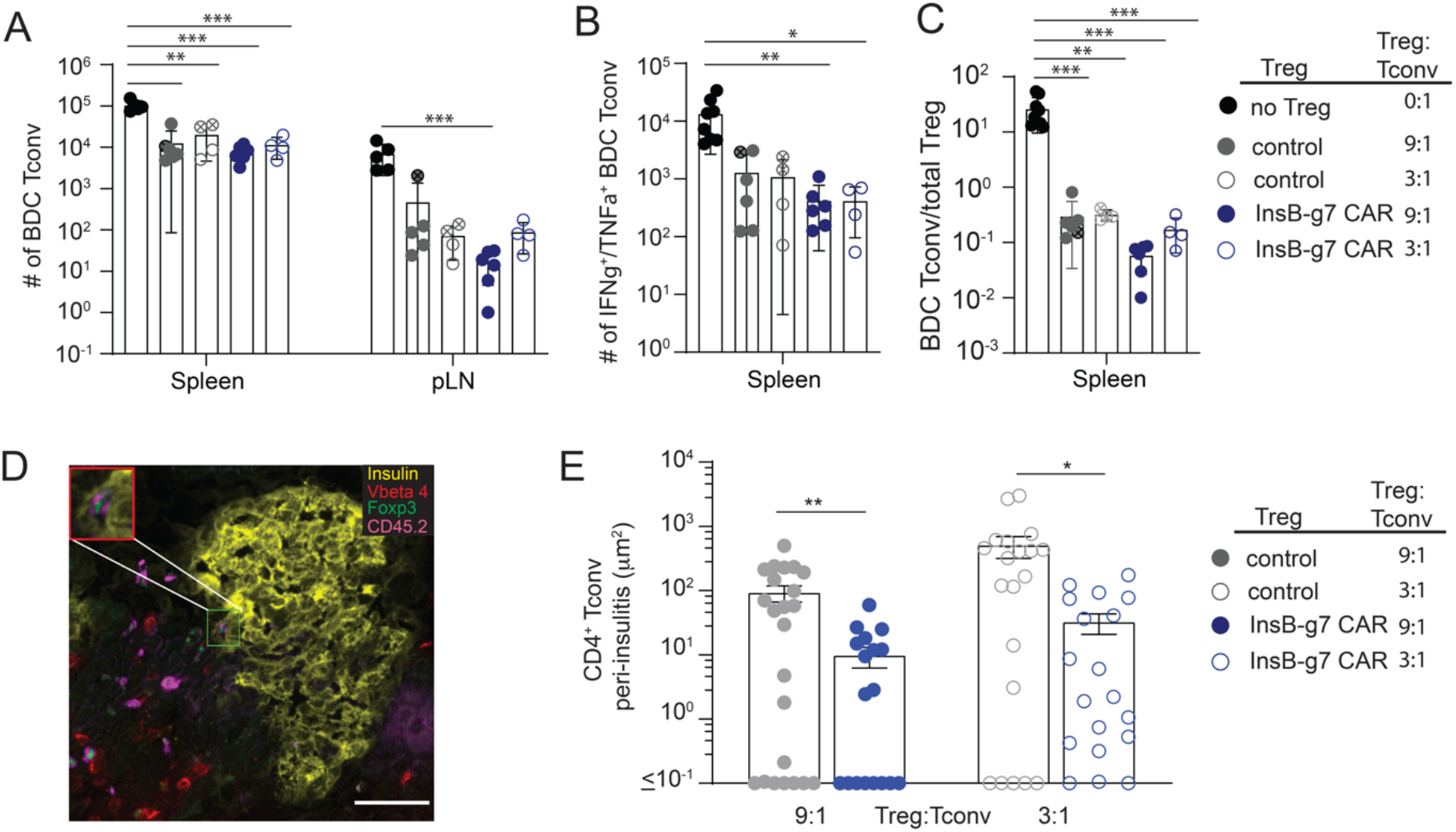
InsB-g7 CAR Tregs reduce the number of BDC 2.5 T effector cells in peripheral lymphoid organs and insulitis. (**A**) Quantification of the number of BDC2.5 Tconv cells in the spleen and pLN from mice described in Fig. 5 A and B. (**B**) Number of IFNγ^+^TNFα^+^ BDC2.5 Tconv cells from spleen of mice described in Fig. 5 A and B and treated at 9:1 or 3:1 Treg:Tconv ratio. (**C**) Ratio of CD4^+^Thy1.1^+^Foxp3^neg^ BDC2.5 Tconv/CD4^+^CD45.2^+^Foxp3^+^Treg from mice described in Fig. 5 A and B. Data are from 2-3 independent experiments, n=4-14 mice/group. One-way ANOVA with multiple comparisons, \**P*<0.05, \*\**P*<0.01, \*\*\**P*<0.001. (**D**) Representative epifluorescence image of a histological section of the pancreas from a mouse described in Fig. 5 A and B at day 30 post-treatment with InsB-g7 CAR Tregs at 3:1 Treg:Tconv ratio. Scale bar indicates 50μm. Inset shows CD45.2^+^(magenta), Foxp3^+^(green) InsB-g7 CAR Tregs and TCR Vβ4^+^(red) BDC2.5 T cells within the peri-islet cellular infiltrate in the pancreas. (**E**) Quantification of total area occupied by Foxp3^−^CD45.2^−^TCR Vβ4^+^CD4^+^ Tconv cells within the peri-islet infiltrate of mice shown in (**B**) analyzed either 2 days after becoming diabetic or at day 30 post-transfer. Data are from 3-4mice/group with 6-10 images/mouse and 88 total images. Kruskal-Wallis test, \**P*<0.05,\*\**P*<0.01.

### InsB-g7 CAR Tregs prevent spontaneous autoimmune diabetes

We next treated wild type NOD mice with InsB-g7 CAR Tregs to determine effects on prevention of spontaneous autoimmune diabetes in an immunocompetent model. InsB-g7 CAR Tregs or vehicle control were transferred into nondiabetic 8-10wk old NOD female mice (Figure 7A). At this age, most female NOD mice have autoantibodies and significant insulitis, similar to stage 1 disease in humans (30). Control mice that received vehicle alone developed diabetes at rates consistent with our colony’s historical incidence, with ∼80% diabetic at 30 weeks of age. In contrast, mice treated with a single injection of InsB-g7 CAR Tregs were significantly protected from diabetes with only 3/7 mice (43%) developing disease by 30 weeks of age (Figure 7B). Despite the significant reduction in autoimmune disease incidence in InsB-g7 CAR Treg-treated mice, at the experimental endpoint of 30 weeks, we could not detect donor CD45.2^+^ cells in the secondary lymphoid organs (including pLN) in either diabetic or non-diabetic mice (data not shown).

**Figure 7.**
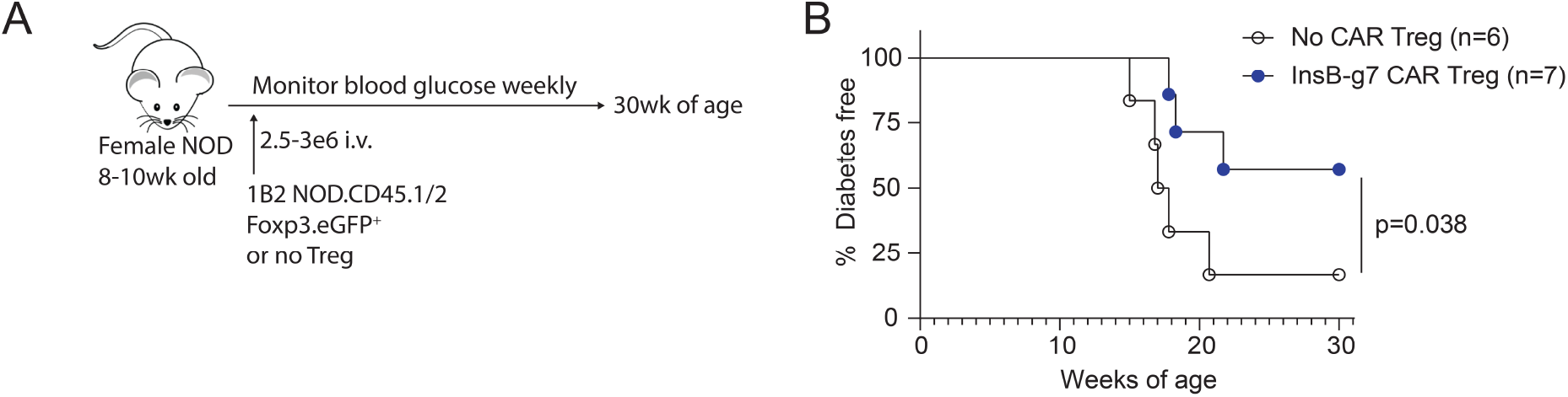
InsB-g7 CAR Tregs prevent spontaneous autoimmune diabetes. (**A**) Experimental design where 8-10 week old NOD female mice received 2.5-3×10^6^ InsB-g7 CAR Tregs or no cells and were monitored until 30 weeks of age for the development of spontaneous diabetes. (**B**) Spontaneous diabetes free survival of mice described in (**A**). Data are pooled from 4 independent experiments, n=6-7mice/group. Gehan-Breslow-Wilcoxon test, \**P*<0.05.

## Discussion

In this study we show that the specificity of Tregs can be redirected towards an islet-derived antigen through the expression of a TCR-like, peptide:MHC Class II specific CAR. Expression of the resulting islet-antigen-specific CAR in Tregs led to enhanced proliferation in response to islet-derived antigen and suppression of effector T cell proliferation and cytokine production *in vitro. In vivo*, CAR Tregs preferentially accumulated in the pancreas-draining lymph nodes where they upregulated molecules associated with activation, resulting in the suppression of effector T cell proliferation, cytokine production and, ultimately, autoimmune diabetes. The potential of TCR-like CARs specific for peptides presented on MHC Class I is increasingly appreciated in cancer (40, 41). To the best of our knowledge this is the first example of the successful use of TCR-like CAR specific for a peptide-MHC Class II complex to control an autoimmune process.

There are a few recent reports on the use of CAR T cells for the treatment of autoimmune diabetes. In one study, an antibody (mAb287) with specificity similar to 1B2 was used to generate a TCR-like CAR and re-direct CD8^+^ T cells to kill APCs presenting InsB in the context of IA^g7^ (42). Although these CD8^+^ CAR T cells delayed diabetes onset in NOD mice, protection declined with time and no significant difference in the overall incidence was observed by 30 weeks of age. Since many different APCs and islet-derived antigens are implicated in autoimmune pathogenesis (43-45), eliminating APCs presenting only one antigen may not be sufficiently potent to prevent activation of diabetogenic T cells. In terms of CAR-Tregs, enforced Foxp3 expression in Tconv cells or co-expression of an anti-insulin scFv-CAR (25) has been explored. Although the anti-insulin CAR Tregs were suppressive *in vitro*, they did not prevent spontaneous autoimmune diabetes in NOD mice, possibly because of low CAR avidity for antigen.

Considering evidence for lack of single-peptide specificity among TCRs (46), we reasoned that an advantage of using a TCR-like scFv CAR rather than a TCR could be increased specificity and affinity. How TCR and CAR affinity/avidity impacts Treg activity remains an open question. There are varying reports of Tregs engineered to express islet-antigen specific TCRs. Human TCR Tregs specific for InsB_11-30_ in the context of HLA-DR3 were significantly less suppressive than influenza hemagglutinin-specific Tregs, but the suppression could be enhanced either through modification of islet-antigen presentation or use of higher affinity mimotopes (47). When TCR Tregs with shared specificity for glutamic acid decarboxylase were compared, cells expressing TCRs with relatively high affinity were more suppressive than lower affinity cells (48). Furthermore, TCR transgenic BDC2.5 Treg cells are potently suppressive *in vitro* and *in vivo* and have high 2D affinity (1.8×10^−3^μM^4^) and functional avidity greater than the well characterized SMARTA CD4 T cells which are specific for foreign antigen gp_66-77_:IA^b^ (7.3×10^−4^μM^4^) (15, 16, 49). In contrast, when Tconv cells were engineered to express FOXP3 and TCRs of varying avidity for IGRP and preproinsulin, functional avidity negatively correlated with suppression (14). Given the lack of consensus on how antigen receptor affinity relates to Treg suppressive function, an interesting future direction would be to compare the functional effects of modified 1B2 scFv CAR with varying affinities, as well as CARs with modified intracellular signalling domains predicted to mediate stronger versus weaker activation signals (50).

Past studies of Tregs in models of autoimmune diabetes showed that antigen-specific Tregs were more suppressive than polyclonal Tregs (13-16, 23, 51, 52). Many of these studies required high numbers (0.1-3×10^6^) of antigen-specific Tregs to inhibit or reverse disease. In WT NOD mice 1.5-2×10^6^ BDC2.5 Tregs were required to prevent islet graft rejection and spontaneous autoimmune diabetes, whereas 5×10^6^ polyclonal Tregs had no effect (15). The numbers of InsB-g7 CAR Tregs used in our study to prevent spontaneous autoimmune disease are in line with these previous studies. BDC2.5 Treg cells can prevent BDC2.5 T cell induced diabetes in NOD.RAGKO mice at a low Treg:Tcell ratio of 1:1, whereas in our study and another report using BDC2.5 Tconv cells engineered to express Foxp3, a minimum ratio of 2:1 TCR or CAR Treg:BDC2.5 T cell ratio was required to prevent disease in immunodeficient NOD mice (14, 15). These data suggest that CAR or TCR Treg therapeutic efficacy can be influenced by TCR or CAR affinity for antigen and/or the context of antigen presentation.

Tregs mediate suppression by a number of mechanisms including production of inhibitory cytokines, metabolic disruption, expression of co-inhibitory molecules, modulating APC function, competition for antigen or cytokines, or direct cytolysis. Evidence that InsB-g7 CAR Tregs potently suppress IL-2 production suggests that consumption of IL-2 could contribute to a negative feedback loop to prevent autoimmunity (53). Further, antigen stimulation of InsB-g7 CAR Tregs resulted in elevated expression of CTLA-4 and significantly reduced CD80 and CD86 expression on the surface of DCs, indicative of transendocytosis (39, 54, 55). Combined with evidence that InsB-g7 CAR Tregs suppress the proliferation and effector function of BCD2.5 T cells *in vitro* and *in vivo*, we speculate that InsB-g7 CAR Tregs function by diminishing costimulation from APCs presenting InsB_10-23_ antigen, thus resulting in bystander suppression of T cells with specificity for disparate islet antigens. Similar bystander suppressive function has recently been reported with TCR-engineered Tregs (14). If modulation of APC function through direct cell contact is a dominant mechanism of Treg-mediated suppression, this would favor the development of Treg therapies that target antigens on the surface of APCs rather than soluble antigens.

Treg homing to target tissue and draining lymph node is required for optimal suppressive function (26). Foxp3^+^ insulin-specific CAR Tregs were detected in the spleen of NOD mice 4 months after a single injection of 2.5×10^6^ cells (25). In another study, DC-expanded BDC2.5 Tregs reversed autoimmune diabetes in NOD mice transplanted with islets, yet donor Tregs could not be detected in pLN or pancreas 50 days after the first of 2 injections. Similarly, we could not find donor Tregs in the secondary lymphoid organs or pancreas of 30 week old NOD mice, nearly 5 months post-injection. Although CXCL10 produced in inflamed islets promotes the recruitment of CXCR3 expressing T cells, this cytokine network is not specific to the islets of Langerhans (56, 57). It could be that transient Treg-mediated suppression at the sites of antigen presentation in pLN nodes may be sufficient to prevent autoimmunity, but in cases where there is ongoing inflammation, it may be necessary for Tregs to access the pancreas.

In conclusion, here we have demonstrated the feasibility of using an InsB_10-23_-IA^g7^ specific mAb to create a peptide-MHC Class II specific CAR. This work provides the first proof-of-concept that CAR Tregs have the potential to be used therapeutically in the context of T1D and in other organ-specific autoimmune or inflammatory disorders.

## Methods

### Mice

NOD mice were purchased from Taconic NOD.CD45.2 (014149), NOD.Foxp3^EGFP^ (025097) (58), NOD.BDC2.5 TCR mice (004460), and NOD. Rag1^KO^ (003729) were purchased from The Jackson Laboratory. NOD.CD90.1, were generated by back crossing BALB/cBy-Thy1a congenic onto NOD/ShiLtJ (001976) for 19 generations and maintained at UMN. NOD.8F10 mice were a generous gift from Dr. Emil Unanue (28). NOD.CD45.2^Het^.Foxp3.eGFP mice were generated by crossing NOD.CD45.2 mice with NOD.Foxp3^EGFP^ mice and F1 mice were used as Treg donors for experiments. NOD.CD90.1.BDC2.5 TCR mice were generated by crossing NOD.CD90.1 mice with NOD.BDC2.5 TCR mice to CD90.1 homozygosity. All mice were housed in specific pathogen–free conditions. The Institutional Animal Care and Use Committee of the University of Minnesota and University of British Columbia approved all animal experiments.

### CARs and Retrovirus

To generate the antigen binding domain of the InsB-g7 CAR, the variable regions of the 1B2 mAb heavy and light chains were sequenced from hybridomas generated as previously described (31, 32). The DNA sequences were then converted into the scFv format and cloned into a murine stem cell virus (MSCV)–based retroviral vector where the scFvs were fused to the hinge (derived from mouse CD8), transmembrane (derived from mouse CD28), and intracellular CD28 and CD3ζ signaling domains. Retroviral particles were produced by using the Platinum-E (Plat-E) Retroviral Packaging Cell Line transfected with the pCL-Eco Retrovirus Packaging Vector, according to the manufacturer’s recommendations (Cell Biolabs). Control Tregs were either cells transduced with a Her2-CAR or left untransduced.

### Isolation, retroviral transduction, and expansion of CAR Tregs

Spleens and lymph nodes (popliteal, axillary, mandibular, and mesenteric) were harvested from 8-12-week-old NOD.Foxp3^EGFP^ mice. The organs were dissociated to release single cells, and CD4^+^ cells were magnetically enriched using mouse negative selection CD4^+^ T cell isolation kits (STEMCELL Technologies and Biolegend). Live Tregs were sorted as fixable viability dye^neg^ (Thermo Fisher Scientific and Tonbo Biosciences), CD4^+^ (BD Biosciences and Tonbo Biosciences), CD25^+^ (BioLegend), and GFP^+^ using either a MoFlo Astrios (Beckman Coulter) or FACS Aria II (BD Biosciences) cell sorter. CD25^neg^, GFP^neg^, CD4^+^ Tconv cells were sorted in parallel. Tregs and Tconvs were cultured in ImmunoCult™-XF T Cell Expansion Medium (STEMCELL Technologies) supplemented with 50 μmol/L of β- mercaptoethanol and 100 units/mL of Penicillin/Streptomycin (Thermo Fisher Scientific). Sorted Treg cultures also contained 1000 U/mL of IL-2 (Proleukin) and 50 nmol/L of rapamycin (Sigma-Aldrich), whereas Tconv cultures contained 100 U/mL of IL-2. Tregs and Tconvs were stimulated with mouse T-Activator CD3/CD28 Dynabeads™ (ThermoFisher Scientific) at a bead to cell ratio of 3:1 and 2:1, respectively.

For transduction, 2 or 3 days post stimulation (for Tconvs or Tregs, respectively), retrovirus, lipofectamine™ 2000 (2 µg/mL, ThermoFisher Scientific) and hexadimethrine bromide (Polybrene, 1.6 µg/mL, Sigma) were added and cells were centrifuged for 90 minutes at 805 x g at 32°C. (17). IL-2 and rapamycin (for Tregs) were replenished when cell cultures were split. On day 7, CD3/CD28 Dynabeads™ were magnetically removed and Tregs and Tconvs were rested for 2 days with decreased IL-2 (300 and 30 U/mL respectively) before use for functional *in vitro* assays. Transduction efficiency was assessed on day 5 and/or 7 post cell activation, by cell surface staining using mouse anti-CD4, anti-Myc, and fixable viability dye eF780. Treg purity was assessed by staining with anti-Foxp3 and anti-Helios. Excess Tregs and Tconvs generated during expansion were cryopreserved on Day 7 post sort following bead removal. All T cells were cryopreserved in 90% ImmunoCult™ base media and 10% dimethyl sulfoxide.

### T cell activation and proliferation assays

On day 9 post stimulation, after two days of cell resting, CAR Tregs and Tconvs were collected, counted, and washed prior to processing in preparation for their respective assays. APCs from NOD spleens were obtained by depleting T cells using the Mouse CD90.2 Positive Selection Kit (STEMCELL Technologies) then pulsed with 15 µM of the indicated peptides and co-cultured with Tregs/Tconvs at a 1:1 ratio. Treg cocultures were supplemented with 100 U/mL IL-2. After 24hrs cells were stained with anti-CD4, anti-Myc, anti-CD69, anti-CTLA-4, anti-LAP, and FVD eF780, and the expression of CD69, LAP, and CTLA-4 were assessed by flow cytometry. For proliferation assays, APCs were irradiated by X-ray at 2000 rad then pulsed with 10 µM peptide. Tregs and Tconvs were labelled with Cell Proliferation Dye CPD eF450 (ThermoFisher Scientific) and cocultured at a 1:1 APC-T cell ratio. CAR Tregs also received supplemental 100 U/mL IL-2 one day into incubation. Cells were stained with anti-CD4, anti-Myc, and FVD eF780, and the proliferation of CAR-Tregs, -Tconvs, and responder T cells was assessed by dilution of their respective cell proliferation dye signal.

### Suppression Assays

To measure suppression of DCs, spleens from NOD mice were dissociated and incubated with Spleen Dissociation Media (STEMCELL Technologies) then DCs were isolated using the EasySep™ Mouse CD11c Positive Selection Kit II (STEMCELL Technologies). While adding the CD11c selection antibody cocktail to the cells, an additional CD11c-BV786 antibody (Invitrogen) was added at the same time to allow for flow cytometry analysis the following day. CD11c^+^ DCs were labelled with CPD eF450 and plated in a 96 well U-bottom plate at either 20,000 or 50,000 per well and pulsed with either 10 μM Insulin P8E or HEL peptide. 100,000 CAR or control Tregs were added to the DCs with IL-2 (100U/mL). Cells were stained with anti-Myc, anti-CD86, anti-CD80, anti-CD11c, anti-CD4, and FVD eF780. Expression of CD80 and CD86 on DCs was measured by flow cytometry after 1 or 2 days.

To measure suppression of BDC2.5 T cells, serial dilutions of CPD eF670-labelled CAR or control Tregs were added to wells with 100,000 CD11c^+^ DCs pulsed with 10 nM p63 and 10 μM Insulin P8E peptides. After 24h, CD4+ T cells from BDC2.5 mice were isolated using the EasySep™ Mouse CD4+ T Cell Isolation Kit, labelled with CPD eF450, and 50,000 were added to wells with Tregs and DCs. After an additional 3 days BDC2.5 T cell proliferation was assessed by flow cytometry and supernatants were collected for analysis with the mouse Th1/Th2/Th17 Cytokine Kit (BD Biosciences) and FCAP Array Software v. 3.0.1 (Soft Flow).

### In vivo experiments

On day 7 post stimulation (4 days after retroviral transduction) Dynabeads were removed and cells were washed and resuspended in Hanks Balanced Salt Solution. CD4^+^ T cells were isolated from NOD.BDC2.5 TCR mice by negative magnetic enrichment using biotinylated antibodies directed against TER119, CD8α, CD11b, CD16/32, NK1.1, Gr-1. Ly6G, B220 (Tobo Biosciences), and MojoSort streptavidin microbeads (BioLegend). Fifty thousand CD4+ BDC2.5 T cells and either 150,000 or 450,000 CAR Treg cells were transferred on the same day via consecutive intravaneous injection into 6-10 week old female NOD.RAG^KO^ mice. Blood glucose was monitored daily starting 7 days post-cell transfer and mice with glucose >250mg/dL (13.9mM) for two or more consecutive days were considered diabetic.

For prevention of spontaneous autoimmune diabetes, 8-10 week old female NOD mice were intravenously injected with 2.5-3 million CD45.2^+^CD4^+^ InsB-g7 CAR Tregs. 1B2 CAR Treg cells were >90% Foxp3^+^ and 70-90% CAR^+^, as indicated by InsBP8E tetramer staining. Littermate cohoused mice injected with Hanks balanced salt solution were used as controls. Blood glucose was monitored once per week until 30 weeks of age with glucose >250mg/dL (13.9mM) for two or more consecutive days were considered diabetic.

### Tetramers and Flow cytometry

To evaluate CAR specificity, cell were stained with various peptide:IA^g7^ tetramers. Soluble peptide:IA^g7^ proteins were produced using S2 insect cells as previously described and made into tetramers by conjugating the soluble peptide:IA^g7^ molecules with BV421 (BioLegend), PE, or APC (Prozyme/Agilent) streptavidin at a 4.5:1 ratio (59). The tetramers used were Insulin B10-23P8E:IA^g7^, p63:IA^g7^, and HEL_11-25_:IA^g7^.

For *in vivo* experiments, single-cell suspensions were obtained from spleen and lymph nodes by mechanical disruption. Lymphocytes were isolated from pancreas by collagenase P and DNase digestion, followed by Percoll density gradient centrifugation. Single cell suspensions were stained with InsBP8E:IA^g7^ tetramers, antibodies against CD4, CD8a, CD11c, CD11b, F4/80, CD90.1, CD90.2, CD45.1, CD45.2, PD-1, and fixable live/dead dye for 30min at 4°C in the presence of Fc block (2.4G2; Bio X Cell), fixed and permeabilized (Tonbo Biosciences, San Diego, CA), and stained for intracellular antigens Foxp3, Helios, IFNγ, TNFα, CTLA-4, and Ki67 overnight at 4°C in permeabilization buffer. A complete list of antibodies can be found in Supplemental Table 1. Flow cytometry was performed on LSRII or Fortessa cytometers (BD Biosciences) and analyzed using FlowJo software v. 10.8 (BD Biosciences).

### Epi-fluorescent microscopy

Pancreata were harvested and frozen in OCT compound (Sakura Finetek) as previously described (60), cut into 4 groups of 8 sequential 7-μm-thick sections separated by a depth of 150-μm using a Leica CM1860 UV cryostat (Leica Microsystems), and mounted as duplicates on Fisherbrand ProbeOn Plus slides glass slides (Thermo Fisher Scientific). Slides were fixed in cold acetone (L10407-AU; Thermo Fisher Scientific) for 10 min and stored at -20°C for no more than 3 months. For imaging, slides were first warmed to 25°C and sections were hydrated in PBS for 10 min. Sections were then blocked with 5% bovine serum albumin (BSA, 9048-46-8; Sigma-Aldrich) in the presence of 1.67μg/mL Fc block (2.4G2; Bio X Cell) in PBS for 1 h at 25°C followed by permeabilization in 5% BSA in the presence of 0.05% Tween-20 (Thermo Fisher Scientific) for 15 minutes at 25°C. Sections were then stained with guinea pig anti-insulin (A0564; Dako, Carpinteria, CA) at 1:1000 and anti-Foxp3 (Alexa Fluor 488, FJK-16s; Thermo Fisher Scientific) at 1:100 overnight at 4°C in 5% BSA in the presence of 0.05% sodium azide (BP922I-500; Thermo Fisher Scientific), 0.5% Triton X-100 (161-0407; Bio-Rad Laboratories), and 1.67ug/mL FC block. Pancreas sections were then stained with secondary and direct conjugate antibodies for 1 h at 25°C in 5% BSA. The secondary antibody used was donkey anti-guinea pig IgG (H and L chain) (Alexa Fluor 647, 706605148; Jackson ImmunoResearch) at 1:1000. Directly conjugate antibodies used at 1:100 included anti-CD45.2 (BV421, 104; BioLegend), anti-CD4 (Alexa Fluor 594, GK1.5; BioLegend), anti-CD8a (PE, 53-6.7; BD Biosciences), and anti-Vβ4 (PE, KT4; BD Biosciences). Slides were mounted with ProlongDiamond antifade reagent (P36961; Thermo Fisher Scientific) using Gold Seal coverslips (12-518-108A; Thermo Fisher Scientific). Images were acquired on a Leica DM6000B epifluorescent microscope with a 20 × objective and quantified using a custom-built Macro (61).

### Statistics

The figures were generated using Adobe Illustrator v. 26 and Graph Pad Prism v. 9.4 (Graph Pad Software, LLC). Statistics tests applied to the data are listed in the figure legends and were calculated using Graph Pad Prism.

### Study Approval

All animal experiments were approved by the Institutional Animal Care and Use Committee of the University of Minnesota (2001-37804A) or University of British Columbia Animal Care Committee (A20-0017).

## Author Contributions

JAS, VF, CMW, MM, PCO, CBV, BTF and MKL designed the study. JAS, VF, CMW, MHA, YC, LAS, AJD, MEW, JSM, MM, PCO, and BTF performed the experiments. NS, MEW and MHA provided animal husbandry. JAS, VF, CMW, MHA, AJD, MEW, JSM, BTF and MKL analyzed the data. JAS prepared the figures. JAS, BTF, and MKL wrote the manuscript. All authors edited the manuscript. JAS and VF are co-first authors, the order reflects the relative contribution to the design and execution of the *in vivo* experiments. BTF and MKL are co-senior authors on the basis of the relative contribution to the initial conception and design of the study.

## Acknowledgments

We thank Drs. Benjamin Williams and Maryaline Coffre from the Helmsley Charitable Trust for their ongoing support and discussion which enabled this work. This work was supported by the resources and staff at the University of Minnesota Center for Flow Cytometry Core Facility with special thanks to Jason Motl, Rashi Arora, and Therese Martin for cell sorting. We acknowledge the Center for Immunology Imaging Core for equipment use and the immunofluorescent image acquisition and analysis directed by Drs. Brian Fife and Jason Mitchell. We thank Jose Nieto from the University of Minnesota Research Animal Resources for animal husbandry. We thank Dr. Lixin Xu for flow cytometry support and the staff at the BCCHR Animal Care Facility for animal husbandry.

## References

1. Brunkow, M.E., et al., Disruption of a new forkhead/winged-helix protein, scurfin, results in the fatal lymphoproliferative disorder of the scurfy mouse. Nat Genet, 2001. 27(1): p. 68–73.

2. Wildin, R.S., et al., X-linked neonatal diabetes mellitus, enteropathy and endocrinopathy syndrome is the human equivalent of mouse scurfy. Nat Genet, 2001. 27(1): p. 18–20.

3. Bennett, C.L., et al., The immune dysregulation, polyendocrinopathy, enteropathy, X-linked syndrome (IPEX) is caused by mutations of FOXP3. Nat Genet, 2001. 27(1): p. 20–1.

4. Brusko, T.M., et al., Functional defects and the influence of age on the frequency of CD4+ CD25+ T-cells in type 1 diabetes. Diabetes, 2005. 54(5): p. 1407–14.

5. Lindley, S., et al., Defective suppressor function in CD4(+)CD25(+) T-cells from patients with type 1 diabetes. Diabetes, 2005. 54(1): p. 92–9.

6. Pesenacker, A.M., et al., A Regulatory T-Cell Gene Signature Is a Specific and Sensitive Biomarker to Identify Children With New-Onset Type 1 Diabetes. Diabetes, 2016. 65(4): p. 1031–9.

7. Pesenacker, A.M., et al., Treg gene signatures predict and measure type 1 diabetes trajectory. JCI Insight, 2019. 4(6).

8. Bluestone, J.A., J.H. Buckner, and K.C. Herold, Immunotherapy: Building a bridge to a cure for type 1 diabetes. Science, 2021. 373(6554): p. 510–516.

9. Marek-Trzonkowska, N., et al., Administration of CD4+CD25highCD127-regulatory T cells preserves beta-cell function in type 1 diabetes in children. Diabetes Care, 2012. 35(9): p. 1817–20.

10. Bluestone, J.A., et al., Type 1 diabetes immunotherapy using polyclonal regulatory T cells. Sci Transl Med, 2015. 7(315): p. 315ra189.

11. Dong, S., et al., The effect of low-dose IL-2 and Treg adoptive cell therapy in patients with type 1 diabetes. JCI Insight, 2021. 6(18).

12. Gitelman, S.E. and J.A. Bluestone, Regulatory T cell therapy for type 1 diabetes: May the force be with you. J Autoimmun, 2016. 71: p. 78–87.

13. Tarbell, K.V., et al., CD25+ CD4+ T cells, expanded with dendritic cells presenting a single autoantigenic peptide, suppress autoimmune diabetes. J Exp Med, 2004. 199(11): p. 1467–77.

14. Yang, S.J., et al., Pancreatic islet-specific engineered T(regs) exhibit robust antigen-specific and bystander immune suppression in type 1 diabetes models. Sci Transl Med, 2022. 14(665): p. eabn1716.

15. Tang, Q., et al., In vitro-expanded antigen-specific regulatory T cells suppress autoimmune diabetes. J Exp Med, 2004. 199(11): p. 1455–65.

16. Tarbell, K.V., et al., Dendritic cell-expanded, islet-specific CD4+ CD25+ CD62L+ regulatory T cells restore normoglycemia in diabetic NOD mice. J Exp Med, 2007. 204(1): p. 191–201.

17. MacDonald, K.G., et al., Alloantigen-specific regulatory T cells generated with a chimeric antigen receptor. J Clin Invest, 2016. 126(4): p. 1413–24.

18. Dawson, N.A., et al., Systematic testing and specificity mapping of alloantigen-specific chimeric antigen receptors in regulatory T cells. JCI Insight, 2019. 4(6).

19. Sicard, A., et al., Donor-specific chimeric antigen receptor Tregs limit rejection in naive but not sensitized allograft recipients. Am J Transplant, 2020. 20(6): p. 1562–1573.

20. Noyan, F., et al., Prevention of Allograft Rejection by Use of Regulatory T Cells With an MHC-Specific Chimeric Antigen Receptor. Am J Transplant, 2017. 17(4): p. 917–930.

21. Boardman, D.A., et al., Expression of a Chimeric Antigen Receptor Specific for Donor HLA Class I Enhances the Potency of Human Regulatory T Cells in Preventing Human Skin Transplant Rejection. Am J Transplant, 2017. 17(4): p. 931–943.

22. Fransson, M., et al., CAR/FoxP3-engineered T regulatory cells target the CNS and suppress EAE upon intranasal delivery. J Neuroinflammation, 2012. 9: p. 112.

23. Blat, D., et al., Suppression of murine colitis and its associated cancer by carcinoembryonic antigen-specific regulatory T cells. Mol Ther, 2014. 22(5): p. 1018–28.

24. De Paula Pohl, A., et al., Engineered regulatory T cells expressing myelin-specific chimeric antigen receptors suppress EAE progression. Cell Immunol, 2020. 358: p. 104222.

25. Tenspolde, M., et al., Regulatory T cells engineered with a novel insulin-specific chimeric antigen receptor as a candidate immunotherapy for type 1 diabetes. J Autoimmun, 2019. 103: p. 102289.

26. Huang, M.T., et al., Lymph node trafficking of regulatory T cells is prerequisite for immune suppression. J Leukoc Biol, 2016. 99(4): p. 561–8.

27. Burton, A.R., et al., On the pathogenicity of autoantigen-specific T-cell receptors. Diabetes, 2008. 57(5): p. 1321–30.

28. Mohan, J.F., et al., Pathogenic CD4(+) T cells recognizing an unstable peptide of insulin are directly recruited into islets bypassing local lymph nodes. J Exp Med, 2013. 210(11): p. 2403–14.

29. Bettini, M.L. and M. Bettini, Understanding Autoimmune Diabetes through the Prism of the Tri-Molecular Complex. Front Endocrinol (Lausanne), 2017. 8: p. 351.

30. Nakayama, M., et al., Prime role for an insulin epitope in the development of type 1 diabetes in NOD mice. Nature, 2005. 435(7039): p. 220–3.

31. Martinov, T., et al., Programmed Death-1 Restrains the Germinal Center in Type 1 Diabetes. J Immunol, 2019. 203(4): p. 844–852.

32. Spanier, J.A., et al., Efficient generation of monoclonal antibodies against peptide in the context of MHCII using magnetic enrichment. Nat Commun, 2016. 7: p. 11804.

33. Stadinski, B.D., et al., Diabetogenic T cells recognize insulin bound to IAg7 in an unexpected, weakly binding register. Proc Natl Acad Sci U S A, 2010. 107(24): p. 10978–83.

34. Crawford, F., et al., Specificity and detection of insulin-reactive CD4+ T cells in type 1 diabetes in the nonobese diabetic (NOD) mouse. Proc Natl Acad Sci U S A, 2011. 108(40): p. 16729–34.

35. Mohan, J.F., S.J. Petzold, and E.R. Unanue, Register shifting of an insulin peptide-MHC complex allows diabetogenic T cells to escape thymic deletion. J Exp Med, 2011. 208(12): p. 2375–83.

36. Levisetti, M.G., et al., The insulin-specific T cells of nonobese diabetic mice recognize a weak MHC-binding segment in more than one form. J Immunol, 2007. 178(10): p. 6051–7.

37. Yamazaki, S., et al., Direct expansion of functional CD25+ CD4+ regulatory T cells by antigen-processing dendritic cells. J Exp Med, 2003. 198(2): p. 235–47.

38. Agata, Y., et al., Expression of the PD-1 antigen on the surface of stimulated mouse T and B lymphocytes. Int Immunol, 1996. 8(5): p. 765–72.

39. Dawson, N.A.J., et al., Functional effects of chimeric antigen receptor co-receptor signaling domains in human regulatory T cells. Sci Transl Med, 2020. 12(557).

40. Li, Y., W. Jiang, and E.D. Mellins, TCR-like antibodies targeting autoantigen-mhc complexes: a mini-review. Front Immunol, 2022. 13: p. 968432.

41. Akatsuka, Y., TCR-Like CAR-T Cells Targeting MHC-Bound Minor Histocompatibility Antigens. Front Immunol, 2020. 11: p. 257.

42. Zhang, L., et al., Chimeric antigen receptor (CAR) T cells targeting a pathogenic MHC class II:peptide complex modulate the progression of autoimmune diabetes. J Autoimmun, 2019. 96: p. 50–58.

43. Ferris, S.T., et al., A minor subset of Batf3-dependent antigen-presenting cells in islets of Langerhans is essential for the development of autoimmune diabetes. Immunity, 2014. 41(4): p. 657–69.

44. Carrero, J.A., et al., Resident macrophages of pancreatic islets have a seminal role in the initiation of autoimmune diabetes of NOD mice. Proc Natl Acad Sci U S A, 2017. 114(48): p. E10418–E10427.

45. Noorchashm, H., et al., B-cells are required for the initiation of insulitis and sialitis in nonobese diabetic mice. Diabetes, 1997. 46(6): p. 941–6.

46. Wooldridge, L., et al., A single autoimmune T cell receptor recognizes more than a million different peptides. J Biol Chem, 2012. 287(2): p. 1168–77.

47. Hull, C.M., et al., Generation of human islet-specific regulatory T cells by TCR gene transfer. J Autoimmun, 2017. 79: p. 63–73.

48. Yeh, W.I., et al., Avidity and Bystander Suppressive Capacity of Human Regulatory T Cells Expressing De Novo Autoreactive T-Cell Receptors in Type 1 Diabetes. Front Immunol, 2017. 8: p. 1313.

49. Liu, B., et al., A Hybrid Insulin Epitope Maintains High 2D Affinity for Diabetogenic T Cells in the Periphery. Diabetes, 2020. 69(3): p. 381–391.

50. Lindner, S.E., et al., Chimeric antigen receptor signaling: Functional consequences and design implications. Sci Adv, 2020. 6(21): p. eaaz3223.

51. Stephens, L.A., K.H. Malpass, and S.M. Anderton, Curing CNS autoimmune disease with myelin-reactive Foxp3+ Treg. Eur J Immunol, 2009. 39(4): p. 1108–17.

52. Elinav, E., T. Waks, and Z. Eshhar, Redirection of regulatory T cells with predetermined specificity for the treatment of experimental colitis in mice. Gastroenterology, 2008. 134(7): p. 2014–24.

53. Liu, Z., et al., Immune homeostasis enforced by co-localized effector and regulatory T cells. Nature, 2015. 528(7581): p. 225–30.

54. Hou, T.Z., et al., A transendocytosis model of CTLA-4 function predicts its suppressive behavior on regulatory T cells. J Immunol, 2015. 194(5): p. 2148–59.

55. Ovcinnikovs, V., et al., CTLA-4-mediated transendocytosis of costimulatory molecules primarily targets migratory dendritic cells. Sci Immunol, 2019. 4(35).

56. Osum, K.C., et al., Interferon-gamma drives programmed death-ligand 1 expression on islet beta cells to limit T cell function during autoimmune diabetes. Sci Rep, 2018. 8(1): p. 8295.

57. Frigerio, S., et al., Beta cells are responsible for CXCR3-mediated T-cell infiltration in insulitis. Nat Med, 2002. 8(12): p. 1414–20.

58. Haribhai, D., et al., Regulatory T cells dynamically control the primary immune response to foreign antigen. J Immunol, 2007. 178(5): p. 2961–72.

59. Pauken, K.E., et al., Cutting edge: type 1 diabetes occurs despite robust anergy among endogenous insulin-specific CD4 T cells in NOD mice. J Immunol, 2013. 191(10): p. 4913–7.

60. Fife, B.T., et al., Interactions between PD-1 and PD-L1 promote tolerance by blocking the TCR-induced stop signal. Nat Immunol, 2009. 10(11): p. 1185–92.

61. Dwyer, A.J., et al., Enhanced CD4(+) and CD8(+) T cell infiltrate within convex hull defined pancreatic islet borders as autoimmune diabetes progresses. Sci Rep, 2021. 11(1): p. 17142.

