## Supplemental data for "Insulin B peptide-MHC class II-specific chimeric antigen receptor-Tregs prevent autoimmune diabetes"

A

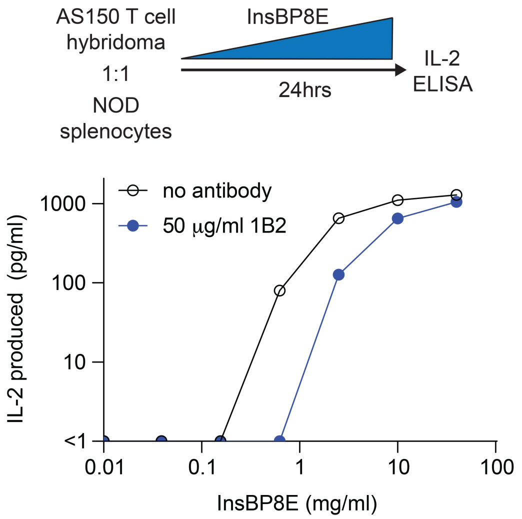


**Supplemental figure 1. 1B2 antibody inhibits IL-2 release by InsBP8E stimulated AS150 T cell hybridoma.** AS150 T cell hybridomas were cultured for 24hrs in the presence of various concentrations of InsBP8E peptide and IL-2 in the hybridoma supernatant was measure by ELISA. Data are representative of two independent experiments, n=1/group.

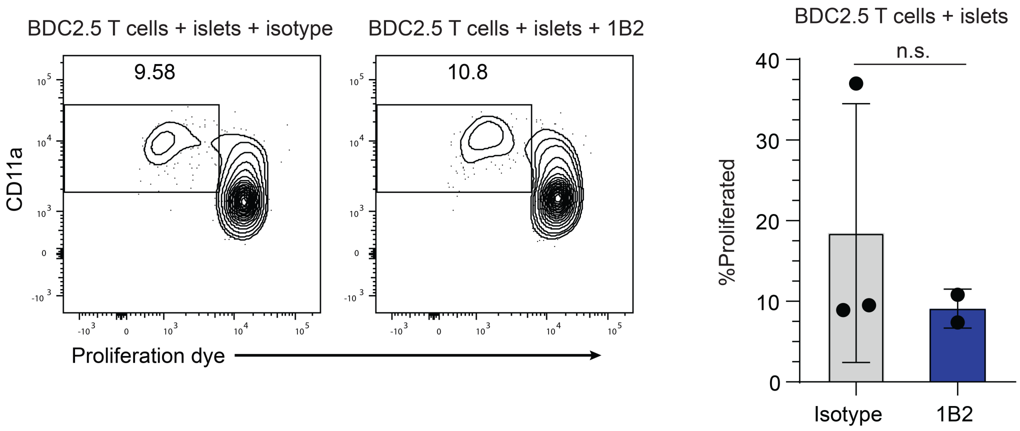


**Supplemental figure 2.** Flow cytometry plots (left) showing CD11a expression, as a marker of activation, and proliferation of BDC2.5 T cells after 3 days in culture with splenocytes as APCs and NOD islets, in the presence of either IgG1 isotype or 1B2 antibody. The panel of the right shows quantification of proliferation from plots shown on the left. Data are representative of 3 independent experiments, n=2-3/group. Students T-test, n.s.=not significant.

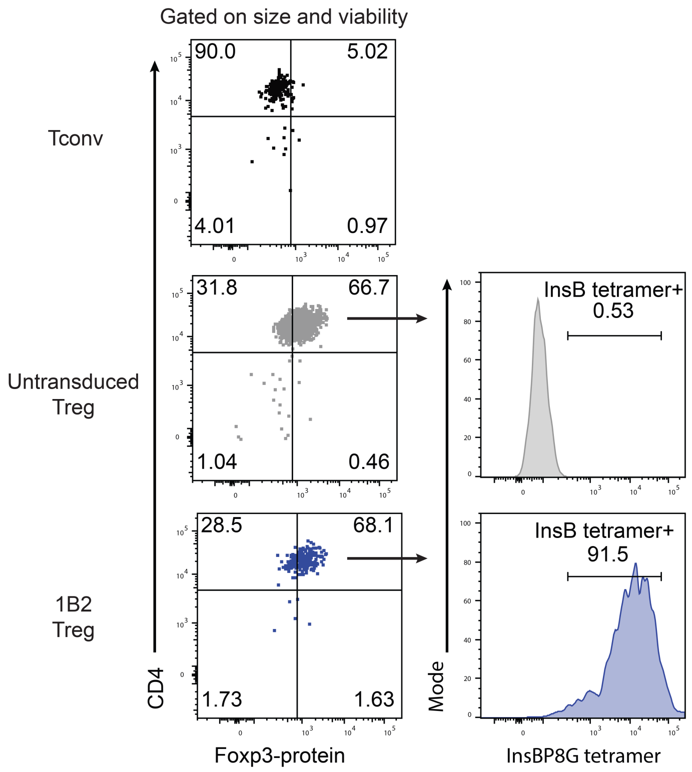


**Supplemental figure 3.** Representative flow cytometry plots of cells after 5 days in culture which were used for *in vivo* experiments. Foxp3 expression was used to determine Treg purity and InsBP8G tetramers were used to determine InsB-g7 CAR retrovirus transduction efficiency.

**Supplemental Table 1**. Antibodies used for flow cytometry

| Antigen | Fluor | Clone | Supplier | Catalogue number |
|---|---|---|---|---|
| CD4 | PE | RM4-5 | Tonbo Biosciences | 50-0042 |
| | | GK1.5 | BD Biosciences | 557308 |
| | BUV496 | GK1.5 | BD Biosciences | 612952 |
| | BV605 | RM4-5 | BD Biosciences | 563151 |
| CD11b | APC-ef780 | M1170 | ThermoFisher | 47-0112-82 |
| CD11c | APC-ef780 | N418 | ThermoFisher | 47-0114-82 |
| | SB780 | N418 | ThermoFisher | 78-0114-80 |
| B220 | APC-ef780 | RA3-6B2 | ThermoFisher | 47-0452-82 |
| Thy1.1 | BUV737 | OX7 | BD Biosciences | 612837 |
| Thy1.2 | BV786 | 30-H12 | BioLegend | 105331 |
| CD45.1 | BV711 | A20 | BioLegend | 110739 |
| CD45.2 | BUV395 | | | |
| PD-1 | PerCp-ef710 | J43 | ThermoFisher | 46-9985-82 |
| Fixable L/D dye | Ghost dye Red-780 | | Tonbo Biosciences | 13-0865 |
| Fixable Viability Dye eFluor™ 780 | eF780 | | ThermoFisher | 65-0865-18 |
| Cell Proliferation Dye eFluor™ 450 | eF450 | | ThermoFisher | 65-0842-90 |
| Cell Proliferation Dye eFluor™ 670 | eF670 | | ThermoFisher | 65-0840-90 |
| Foxp3 | AF488 | FJK-16s | ThermoFisher | 53-5773-82 |
| | PE | FJK-16s | ThermoFisher | 12-5773-82 |
| Ki67 | BUV737 | solA15 | ThermoFisher | 367-5698-82 |
| | BV421 | 16A8 | BioLegend | 652411 |
| CTLA4 | PE | 4C10-4B9 | BioLegend | 563151 |
| Helios | PE-Cy7 | 22F6 | BioLegend | 137236 |
| | eF450 | 22F6 | ThermoFisher | 48-9883-42 |
| IFN$\gamma$ | BV650 | XMG1.2 | BioLegend | 505532 |
| TNF$\alpha$ | Ef450 | mAb11 | ThermoFisher | 48-7349-42 |
| c-Myc | AF647 | 9E10 | UBC Ablab | 67-0029-01 |
| LAP | PE | TW7-16B4 | ThermoFisher | 12-9821-82 |
| CD69 | BV785 | H1.2F3 | Biolegend | 104543 |
| CD152 (CTLA-4) | BV605 | UC10-4B9 | Biolegend | 106323 |
| CD86 | BV650 | GL1 | BD Biosciences | 564200 |
| CD80 | BUV395 | 16-10A1 | BD Biosciences | 740246 |
